# Structure of small HBV surface antigen reveals mechanism of dimer formation

**DOI:** 10.1101/2024.09.13.612767

**Authors:** Xiao He, Yunlu Kang, Weiyu Tao, Jiaxuan Xu, Xiaoyu Liu, Lei Chen

## Abstract

Hepatitis B surface antigen (HBsAg), the only membrane protein on the HBV viral envelope, plays essential roles in HBV assembly, viral release, host cell attachment, and entry. Despite its functional and therapeutic importance, the detailed structure of HBsAg has remained enigmatic. Here, we present the core structure of HBsAg at 3.6 Å resolution, determined using recombinant small spherical subviral particles (SVPs). The structure reveals how two HBsAg monomers interact to form a dimer, which is the basic building block of SVPs.

## Main

Hepatitis B Virus (HBV) infection can lead to chronic hepatitis B, which significantly increases the risk of death from conditions such as cirrhosis and liver cancer ^1^. In 2019, approximately 296 million individuals were living with chronic hepatitis B, and there were about 1.5 million new infections each year ^1^. Moreover, hepatitis B resulted in an estimated 820,000 deaths in 2019, with cirrhosis and hepatocellular carcinoma being the primary causes ^1^. HBV is an enveloped virus with HBsAg as the only protein on its envelope. HBsAg exists in three variants: large (L-HBsAg), medium (M-HBsAg), and small (S-HBsAg) surface antigens, all of which contain the amino acids of S-HBsAg. S-HBsAg is predicted to have four helices (H1-H4) with an extracellular N-terminus ^2^. The cytosolic loop (CYL) is located between H1 and H2, which is important for HBV virion assembly and maturation ^3^. Between H2 and H3, there is a hydrophilic domain known as the “a”-determinant or antigenic loop (AGL), which is essential for the viral infection process ^2^. Unlike S-HBsAg, M-HBsAg contains an additional PreS2 region, while L-HBsAg possesses both PreS1 and PreS2 regions. Infected hepatocytes secrete three forms of HBV-related particles: the 42-nm virions, tubular subviral particles (SVPs), and 22-nm small SVPs ^2^. Small spherical SVPs are protein-lipid complexes formed primarily by S-HBsAg dimers and are the major format used in prophylactic HBV vaccines ^4^. SVPs do not contain viral genomes and are 10^3^-fold to 10^6^-fold more abundant than virions ^2^.

Although the functional significance of HBsAg is well recognized, its structural information is limited to the recently determined medium-resolution structure of small spherical SVP at 6.3 Å ^5^. This structure reveals the overall architecture of the HBsAg dimer and how 24 HBsAg dimers assemble into a rhombicuboctahedral protein complex ^5^. However, the details of how HBsAg monomers interact to form a dimer at the amino acid level and how the CYL contributes to the structural integrity of the S-HBsAg dimer remain unknown. To understand these mechanisms, we sought to determine its structure at high resolution.

Due to limited access to HBV patient serum, we developed a method to recombinantly express the spherical SVP. Given that the N-terminus of mature HBsAg is located extracellularly ^2^, we appended a signal peptide before M-HBsAg constructs (serotype ayw, genotype D3) to direct the insertion of the first transmembrane helix with its N-terminus outside. This approach allowed for the efficient secretion of small spherical SVP into the medium of FreeStyle 293F cells. We further purified these SVPs using a strep-tagged scFV fragment of the NAb HBC34 (referred to as NAb_HBC_) (Fig. S1a). NAb_HBC_ is the parental antibody of the broad NAb VIR-3434, which is in clinical trials for the treatment of HBV and HDV ^6^. NAb_HBC_ recognizes a structural epitope of the AGL of HBsAg ^6^. The purified recombinant SVP was subjected to cryo-EM sample preparation and single-particle analysis (Fig. S1b-c and Fig. S2).

Single-particle reconstruction without symmetry resulted in a map at 4.7 Å resolution with clear helical structural features (Fig. S2). The map shows many protrusions from the near-spherical core, representing the AGL regions (Fig. 1a-d). Further inspection revealed that the recombinant SVP has a pseudo-octahedral symmetry. Three HBsAg dimers are arranged around a three-fold symmetry axis to form the HBsAg hexamer (3×2mer) (Fig. 1a). Seven HBsAg hexamers further assemble into an SVP (Fig. 1). This packing also results in additional two-fold (2×2×2mer) (Fig. 1b) and four-fold (4×2mer) axes (Fig. 1c).

**Fig. 1:**
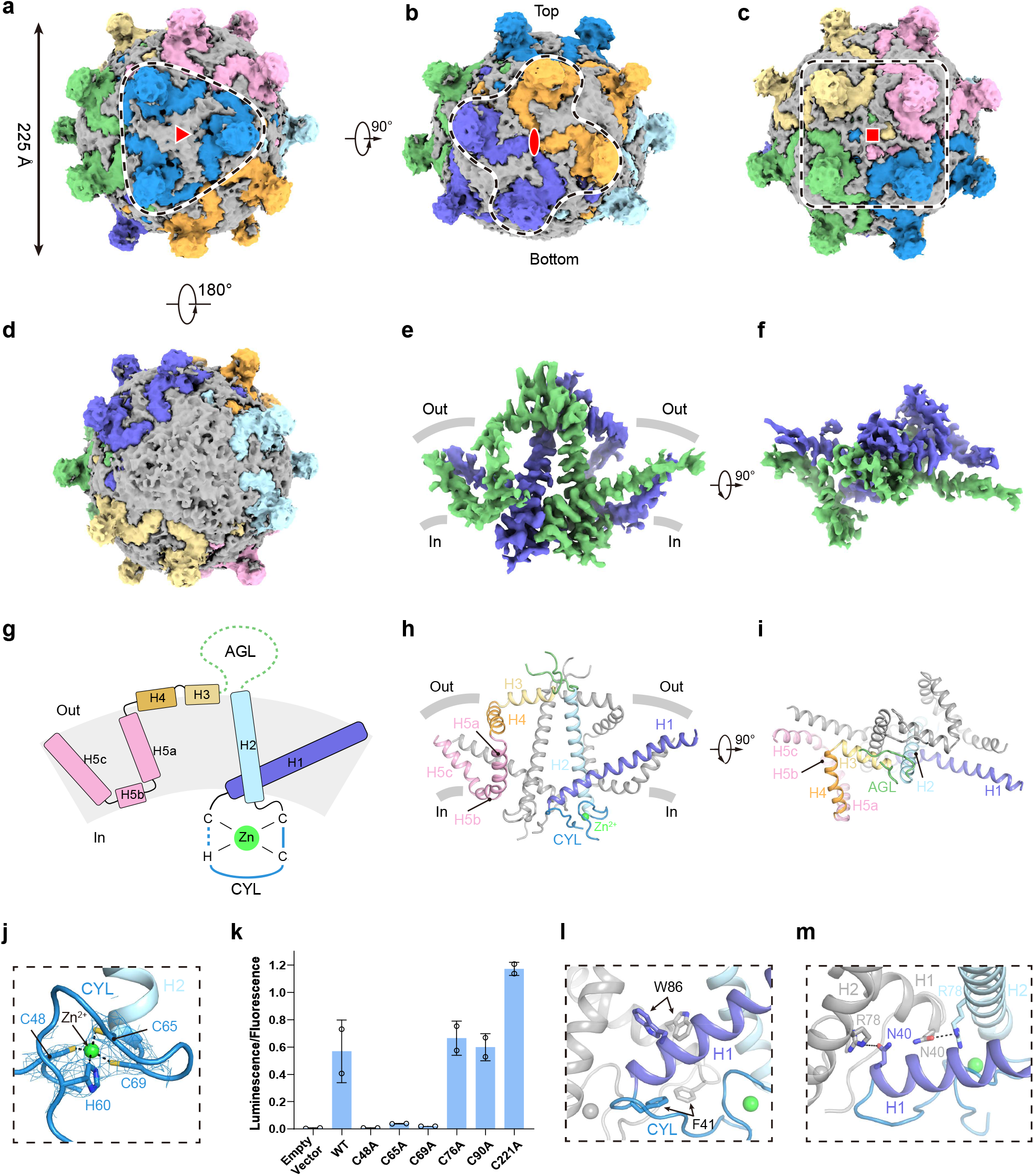
The overall architecture of SVP and structure of HBsAg dimer in SVP. **a**, Top view of of the cryo-EM map of spherical SVP refined in C1 symmetry. Each HBsAg trimer of dimer (3×2mer) in SVP is colored differently, and the putative lipid density is shown in grey. HBsAg 3×2mer is indicated with a dashed line, and the three-fold axis is indicated in its center. **b**, A 90^°^-rotated view of **a**. HBsAg 2×2×2mer is indicated with a dashed line, and the two-fold axis is indicated in its center. Please note the 3×2mer on the top has strong density, while the density of 3×2mer on the bottom is missing. **c**, A 90^°^-rotated view of **a**. HBsAg 4×2mer is indicated with a dashed line, and the four-fold axis is indicated in its center. **d**, A 180^°^-rotated bottom view of **a**. Please note the density of the HBsAg 3×2mer on the bottom is missing. **e**, The side view of the cryo-EM map of HBsAg dimer. Each subunit of the dimer is shown in green (protomer A) or purple (protomer B). The approximate boundaries of the phospholipid bilayer are indicated as thick grey lines. **f**, A 90^°^-rotated top view of **e**. **g**, Topology of one of the protomer of HBsAg dimer. Helices, CYL, and AGL are colored differently, and invisible regions in Cryo-EM map are indicated with dashed lines. **h**, Cartoon representation of HBsAg dimer. Helices, CYL, and AGL in protomer A are colored the same as in g, and protomer B is shown in grey. **i**, A 90^°^-rotated top view of **h**. **j**, Structure of CHC2-type zinc finger in CYL. Zinc ion is shown as bright green sphere, and surrounding cysteines and histidine are shown as sticks. The density of the zinc finger is shown as blue mesh. **k**, Surface expression of various cysteine mutants of GFP-M-HBsAg. n = 2 technical replicates. The experiment was performed three times with similar results. **l**, Hydrophobic interactions between protomers A and B. Key residues are indicated and shown as sticks. **m**, Hydrophilic interactions between protomers A and B. Key residues are indicated and shown as sticks.

However, the electron density is not uniformly strong across the cryo-EM map, suggesting substantial structural heterogeneity (Fig. 1b). We found that HBsAg 3×2mer at the top of the SVP shows strong density (Fig. 1b), while the density of HBsAg 3×2mers between the top and bottom gradually weakens (Fig. 1b), and the density of HBsAg 3×2mers at the bottom is very weak and hardly visible at low contour levels (Fig. 1b and 1d), suggesting that the packing of HBsAg hexamers on SVP is heterogeneous and imperfect compared to the standard octahedral symmetry. This phenomenon was not observed previously in SVPs purified from patient serum, which showed rhombicuboctahedral symmetry ^5^. Whether this discrepancy is due to different expression systems, sample preparation, or single-particle data analysis procedures remains to be elucidated. Because of the deviation from octahedral symmetry in the map (Fig. 1b), forcing reconstruction with octahedral symmetry led to smeared and uninterpretable density maps. Therefore, to improve resolution, we aligned the map to the C3 symmetry axis of the top 3×2mer and subsequent local refinement yielded a consensus map at 4.24 Å resolution (Fig. S2c). To improve the resolution, we further subtracted the 3×2mer, 4×2mer, and 2×2×2mer from the consensus map and refined them individually by applying local symmetry, resulting in maps at 3.5 Å, 3.9 Å, and 3.1 Å resolution, respectively (Fig. S2c). To obtain a better map of the HBsAg dimer, we expanded the HBsAg hexamers (3×2mer) at the top using C3 symmetry and focused the local refinement on a single HBsAg dimer, yielding a map at 3.60 Å averaged resolution (Fig. S2c-h). This map was used for model building of the HBsAg dimer core (Fig. S3 and Table S1). The architecture of the HBsAg dimer core is overall similar to the previously reported medium-resolution V-shaped structure (Fig. 1e-f)^5^, but with enhanced side chain features in many regions (Fig. 1e-f and Fig. S3). Although it was predicted that HBsAg contains four helices ^2^, we found that residues in the last two predicted helices form three short helices (H3-H5) (Fig. 1g-i). Therefore, we have renamed the helices of HBsAg as H1-H5 according to our current structure (Fig. 1g-i and Fig. S4). In addition, H5 is highly curved and is further divided into H5a, H5b and H5c (Fig. 1g-h and Fig. S4). The map quality of the central region of HBsAg is better than other portions (Fig. 1e and Fig. S2h), likely reflecting the remarkable structural dynamic of HBsAg. As a result, we observed discrete side chain density for most of H1, CYL, H2, and H3, but the side chain features for H4-H5 are only barely visible for a few bulky residues (Fig. S3). Additionally, residues 111-149 of the AGL could not be modeled in this map due to the poor local resolution (Fig. 1e-f), indicating its large conformational heterogeneity.

The structure shows that on the extracellular side, the amphipathic helices H3 and H4 of both HBsAg protomers are located on either side of the AGL and float on the viral envelope (Fig. 1g-i). It is previously reported that H3 helix is critical for morphogenesis of SVP^7^. On the intracellular side, instead of being a flexible loop, the CYL forms a defined structure stabilized by a CHC2-type zinc finger structure, with C48, H60, C65, and C69 coordinating a zinc ion (Fig. 1g and j). To investigate the functional importance of the zinc finger motif, we mutated cysteines residues on the zinc finger or in the transmembrane domains of HBsAg into alanines and detect the surface expression of HBsAg. We found that mutations of cysteines on zinc finger (C48A, C65A, and C69A) abolished the surface expression of HBsAg, while cysteines mutants in the transmembrane domain retain robust surface expression (Fig. 1k), suggesting that the integrity of zinc finger is essential for HBsAg maturation. This observation is consistent with previous findings showing that deleting residues in the CYL impair virus production ^3^ and also correlates with the fact that zinc finger-forming residues are absolutely conserved in HBV-related viruses, while cysteines in the transmembrane domain are not conserved (Fig. S4). Interactions between the two HBsAg protomers involve multiple residues at the bottom of H1, CYL, and H2 (Fig. 1l-m). Specifically, W36 on H1 and F41 on CYL interact with their counterparts of the other protomer through hydrophobic interactions (Fig. 1l). Additionally, R78 on H2 makes polar interactions with N40 on H1 of the other protomer (Fig. 1m). These non-covalent interactions stabilize the HBsAg dimer near the intracellular side, complementing the inter-subunit covalent disulfide linkages in the AGL on the extracellular side.

To visualize how HBsAg dimers are assembled into small viral particles (SVP), we docked the HBsAg dimer core structures back into the density map of SVP (Fig. S5). On the three-fold symmetry axis, the upper regions of H1 from three HBsAg dimers pack together to form a three-helical bundle (Fig. S5a). H5c of one dimer interacts with H1 and H2 of adjacent dimers (Fig. S5). On the four-fold axis, the upper regions of H1 from four dimers pack together to form a four-helical bundle, and H5c of one dimer interacts with H1 of an adjacent dimer (Fig. S5b). Close to the two-fold axis, H5a of one dimer interacts with H5a of the other dimer (Fig. S5c). These inter-dimer interactions stabilize the overall packing and assembly of the spherical SVP.

SVP is the major format of the prophylactic HBV vaccines ^4^. The high-resolution structures of the HBsAg dimer core and its higher assembly in the small spherical SVP presented here provide a near-atomic resolution understanding of the assembly mechanism of SVP. This information might offer clues for further drug discovery targeting the folding or maturation of the HBsAg protein or for the structure-based development of next-generation HBV vaccines that are more stable. Furthermore, the highly dynamic nature of H1 and H4-H5 helices of HBsAg observed here might be crucial for the plasticity of HBsAg dimer and their assembly into HBV viral envelope or tubular SVP, which might have different packing environments compared to the spherical SVP.

## Data availability

Cryo-EM maps and the atomic coordinate of the HBsAg-dimer core have been deposited in the EMDB and PDB under the ID codes EMDB: EMD-61003 and PDB: 9IYX, respectively

## Acknowledgements

We thank Prof. Christoph Seeger, Prof. John E. Tavis, Prof. Wang-Shick Ryu, and Prof. Joseph Anderson for providing the HBV DNA. We thank Miao Wei and Rui Liu for helpful suggestions on manuscript writing. Cryo-EM data collection was supported by the Electron microscopy laboratory and the Cryo-EM platform of Peking University with the assistance of Xuemei Li, Zhenxi Guo, Changdong Qin, Xia Pei, Xiaojuan Hui, and Guopeng Wang. Part of the structural computation was also performed on the Computing Platform of the Center for Life Science and High-performance Computing Platform of Peking University. We thank the National Center for Protein Sciences at Peking University in Beijing, China for assistance with negative stain EM. The work is supported by grants from the Ministry of Science and Technology of China (National Key R&D Program of China, 2022YFA0806504 to L.C.), the National Natural Science Foundation of China (31821091 to L.C.), and the Center for Life Sciences (CLS to L.C.).

## Authors contributions

L.C. initiated the project and wrote the manuscript draft. X.H. purified the protein, and prepared the cryo-EM sample. H.X., J.X., and X.L. collected the cryo-EM data. X.H., Y.L., and L.C. processed the data and built the model. H.X., and W.T. made cysteine mutations and purified antibodies. All authors contributed to the manuscript preparation.

## Supplementary Information for

## Methods

### Cell Culture

Sf9 insect cells (Thermo Fisher Scientific) were cultured in SIM SF (Sino Biological) at 27 °C. FreeStyle 293F suspension cells (Thermo Fisher Scientific) were cultured in FreeStyle 293 medium (Thermo Fisher Scientific) supplemented with 1% fetal bovine serum (FBS, VisTech), 67 μg ml^−1^ penicillin (Macklin), and 139 μg ml^−1^ streptomycin (Macklin) at 37°C with 6% CO_2_ and 70% humidity. The cell lines were routinely checked to be negative for mycoplasma contamination but have not been authenticated.

### Expression and purification of small spherical SVP

BacMam virus carrying rSP-MHBsAg was generated with SF9 cells. 293F cells at a density of 2.5×10 ^6^ cells mL^−1^ were transfected with 8% volume of P2 virus. 10 mM sodium butyrate was added 12 h after infection and the cells were incubated at 37 ^°^C for 60h before harvest. After 4000 rpm for 10 min, the cell medium was collected, and the cell debris and insoluble aggregated proteins were removed by 12000 rpm for 15 min (JA 14). To precipitate SVP, 10% PEG6000 (w/v) was added into the collected medium, thoroughly mixed on a magnetic stirrer for 30 min followed by incubation on ice for another 30 min. After 12000 rpm for 15 min (JA 14), the precipitate was collected and resolved with TBS. Insoluble materials were removed by 16000 g for 15 min (JA 14). For purification of SVP, scFv_HBC_-strep was expressed and purified the same as Fab_HBC_, and the supernatant was added with scFv_HBC_-strep and subjected to strep affinity chromatography to obtain the scFv_HBC_-binding SVPs. The scFv_HBC_-bound SVPs were eluted with buffer containing 50 mM HEPES (pH=8), 150 mM NaCl, and 5 mM desthiobiotin, and subjected to cryo-EM sample preparation.

### Cryo-EM sample preparation and data collection

For the SVP sample, the purified scFv_HBC_-SVP was diluted to A_280_ = 2.0. To avoid protein denaturation at the air-water interface, 0.01% glycyrrhizin (estimated concentration of 122 μM) was added to the sample before cryo-EM sample preparation. Holey carbon grids (Quantifoil Au 300 mesh, R 0.6/1) were coated with freshly prepared graphene oxide. Aliquots of 3 μl purified protein were applied on glow-discharged grids and the grids were blotted for 6 s before being plunged into liquid ethane using Vitrobot Mark IV (Thermo Fisher Scientific). Cryo-grids were first screened on a Talos Arctica electron microscope (Thermo Fisher Scientific) operating at 200 kV with a K2 camera (Gatan). The screened grids were subsequently transferred to a Titan Krios electron microscope (Thermo Fisher Scientific) operating at 300 kV with a K3 camera (Gatan) and a GIF Quantum energy filter (Gatan) set to a slit width of 20 eV. Images were automatically collected using EPU (Thermo Fisher Scientific) in super-resolution mode at a nominal magnification of ×81,000, corresponding to a calibrated super-resolution pixel size of 0.533 Å with a preset defocus range from –1.5 to –1.8 μm. Each image was acquired as a 3.74-s movie stack of 47 frames with a dose rate of 21.46 e^−^ Å^−2^ s^−1^, resulting in a total dose of about 70 e^−^ Å^−2^.

### Cryo-EM image analysis

The image processing workflows are illustrated in Supplementary Figures. Super-resolution movie stacks were collected. Motion-correction, two-fold binning, and dose weighting were performed using MotionCor2 ^1^. Contrast transfer function (CTF) parameters were estimated with cryoSPARC ^2^. Micrographs with ice or ethane contamination and empty carbon were removed manually. Particles were auto-picked using Gautomatch (provided by K. Zhang). All subsequent classification and reconstruction were performed in cryoSPARC ^2^ unless otherwise stated. Reference-free 2D classification was performed to remove contaminants. The resulting particles were subjected to 3D classification using the initial models generated from cryoSPARC. For the SVP dataset, extensive 2D and 3D classification resulted in 530,120 particles, which were subjected to ab initio reconstruction and non-uniform refinement using C1 symmetry, resulting in a map at 4.70Å resolution. The map was aligned to the C3 symmetry axis on the top HBsAg hexamer (3×2mer) and subjected to local refinement using C3 symmetry, resulting in a consensus map at 4.24Å resolution. The 3×2mer on top and 2×2×2mer on side was subtracted out of the particles of this consensus refinement. For 4×2mer, the particles of consensus refinement were expended using C3 symmetry and then used for subtraction. The subtracted particles were subjected to local refinement with corresponding symmetry, yielding maps at 3.49Å for 3×2mer with C3 symmetry, 3.40Å for 4×2mer with C4 symmetry, and 3.90 Å for 2×2×2mer with C2 symmetry. To further improve the resolution of the HBsAg dimer, the particles of locally refined 3×2mer were subjected to symmetry expansion using C3 symmetry and signal subtraction to obtain signals of a single HBsAg dimer. The resulting particles were subjected to local refinement, yielding a map at a resolution of 3.60Å.

### Model building

For SVP, the initial model of HBsAg-dimer was predicted with ColabFold ^3^. The transmembrane H1 and H2 were fitted into the maps using UCSF Chimera ^4^ and rebuilt manually in Coot ^5^. Model refinement was performed using phenix.real_space_refine in PHENIX ^7^. The validation statistics are provided in Tables S1 and S2.

### Surface labelling of HbsAg

AD293 cells were seeded on 24-well dish coated with poly-D-lysine. After cells were adherent, wild type GFP-M-HBsAg and the mutants were transfected. For surface labeling, the cells were washed with PBS for two times 36 h after transfection, and the whole experiment was performed at room temperature. Cells were incubated with 4 % formaldehyde (10.8% formalin) for 30 min for fixation, and after washed with 500 uL PBS for two times, fixed cells were incubated with 300 uL blocking buffer (3% goat serum in PBS). After 30 min, Hep B preS2 Antibody (Santa Cruz sc-23944, 1:2500 diluted in blocking buffer) was added to specially combine with HBsAg that had been expressed and transported onto cell membrane eventually. After incubation for 1 h, unbonded antibodies were removed by PBS washing for 3 times, and goat anti-mouse IgG conjugated with HRP (Invitrogen 31444, 1:2500 diluted in blocking buffer) was added and incubated for 1 h. After extensive washing with PBS, the cells were added with 250 uL ECL substrate (Tanon 180-5001, 1:20 diluted in water) and reacted for 90 s. Luminescence and fluorescence intensity (488 nm excitation and 520 nm emission) signals were measured with an Infinite M Plex plate reader (Tecan). For normalize, the values of luminescence were divided with fluorescence intensity values.

### Quantification and statistical analysis

Global resolution estimations of cryo-EM density maps are based on the 0.143 Fourier Shell Correlation criterion ^8^. The local resolution was estimated using cryoSPARC. The number of independent experiments (N) and the relevant statistical parameters for each experiment (such as mean or standard deviation) are described in the figure legends. No statistical methods were used to pre-determine sample sizes.

---

**Fig. S1.**
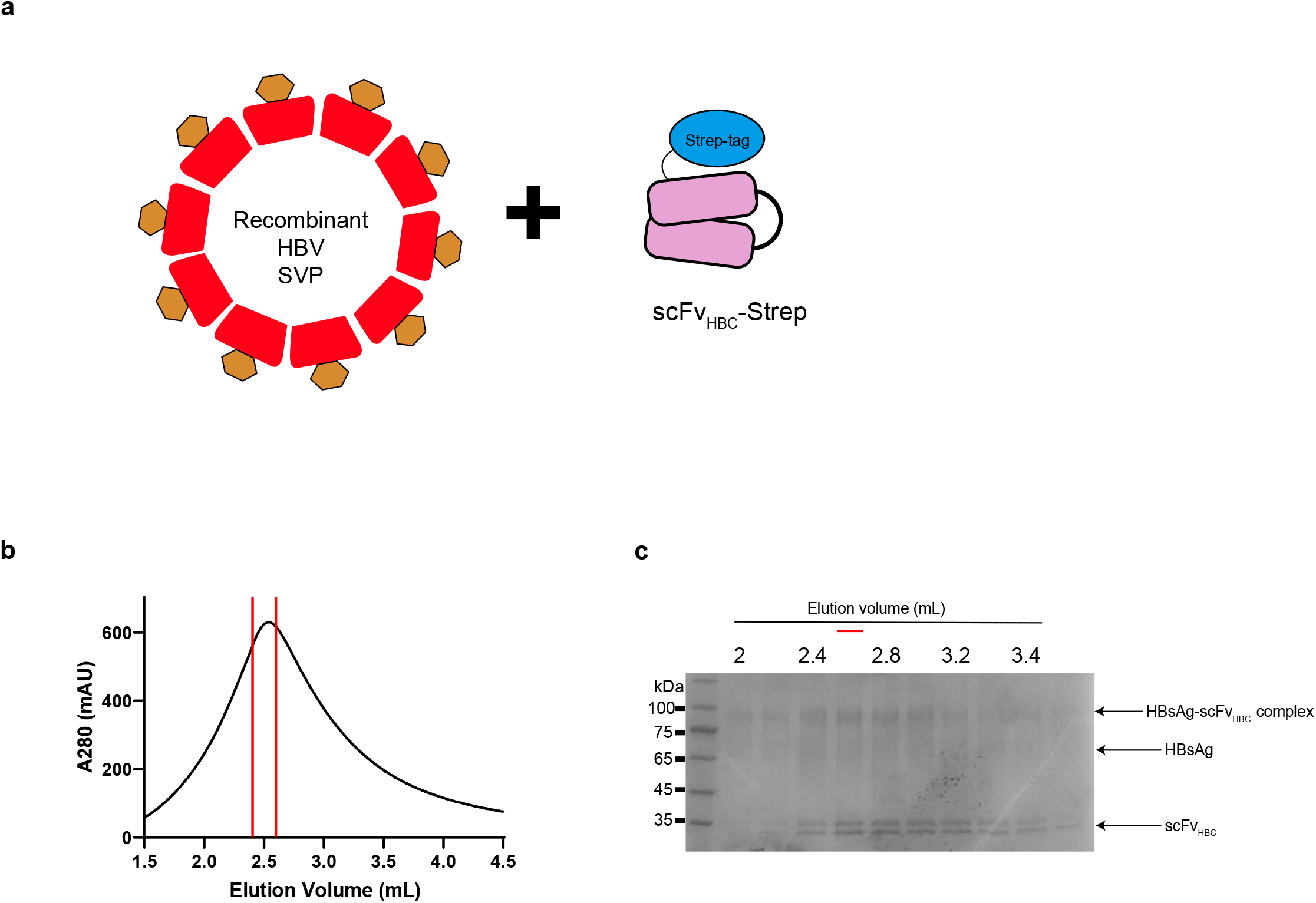
Purification of recombinant small spherical SVP. **a**, Schematic diagram of the strategy of SVP purification using NAb_HBC_. Affinity chromatography was performed by using the strep tag on the scFv fragment. **b**, Strep affinity chromatography of the SVP-scFv_HBC_ complex. The fraction that contains the highest concentration of protein (indicated by red lines) was subjected to cryo-EM sample preparation. **c**, Coomassie brilliant blue staining of SDS-PAGE of fractions from affinity chromatography in **b**. The bands corresponding to HBsAg-scFv_HBC_ complex, HBsAg and scFv_HBC_ were indicated. The fractions indicated by the red line were used for cryo-EM sample preparation.

**Fig. S2.**
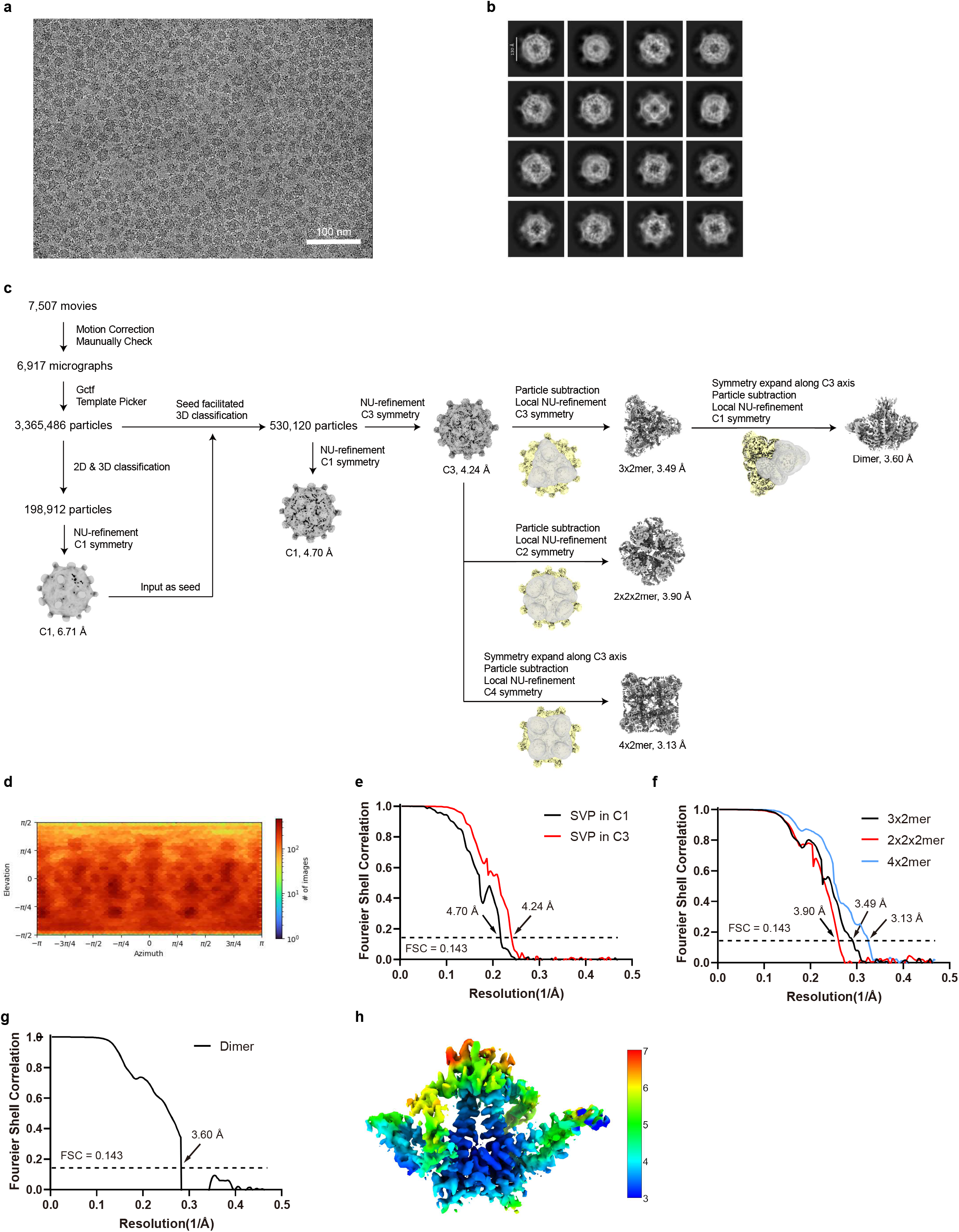
Cryo-EM image analysis of spherical SVP. **a**, Representative raw micrograph (7,507 in total) of spherical SVP. Scale bar, 100 nm. **b**, 2D-class averages of spherical SVP. Scale bar, 130 Å. **c**, Cryo-EM data processing workflow of spherical SVP, 3×2mer, 2×2×2mer, 4×2mer and dimer. For details, see ‘Cryo-EM image analysis’ in the Methods section. **d**, Angular distribution of spherical SVP in C1 symmetry. **e**, Gold standard Fourier Shell Correlation (FSC) of spherical SVP after correction of masking effects. **f**, Gold standard Fourier Shell Correlation (FSC) of HBsAg 3×2mer, 2×2×2mer, and 4×2mer after correction of masking effects. **g**, Gold standard Fourier Shell Correlation (FSC) of HBsAg dimer after correction of masking effects. **h**, Local resolution map of HBsAg dimer. Scale bar, 3–7 Å.

**Fig. S3.**
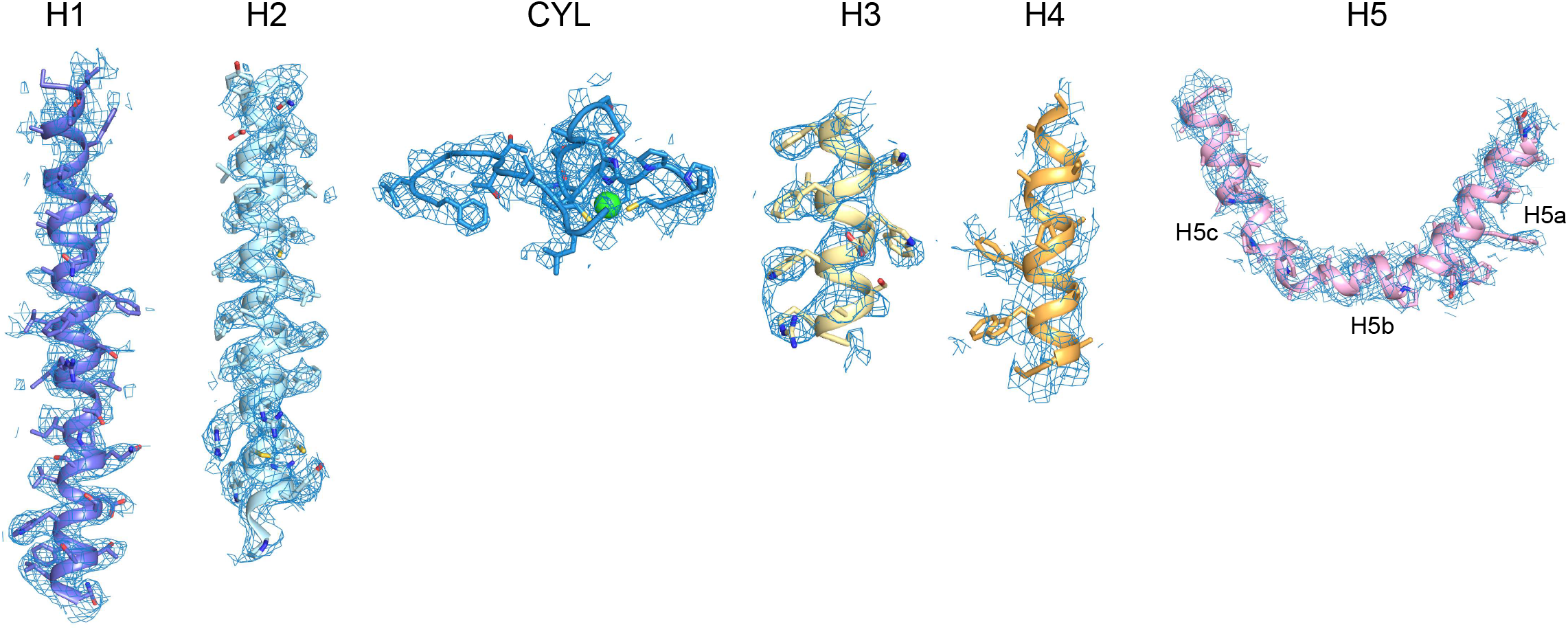
Representative electron density maps of HBsAg dimer. Electron density maps of helices and CYL in HBsAg dimer are shown in blue meshes.

**Fig. S4.**
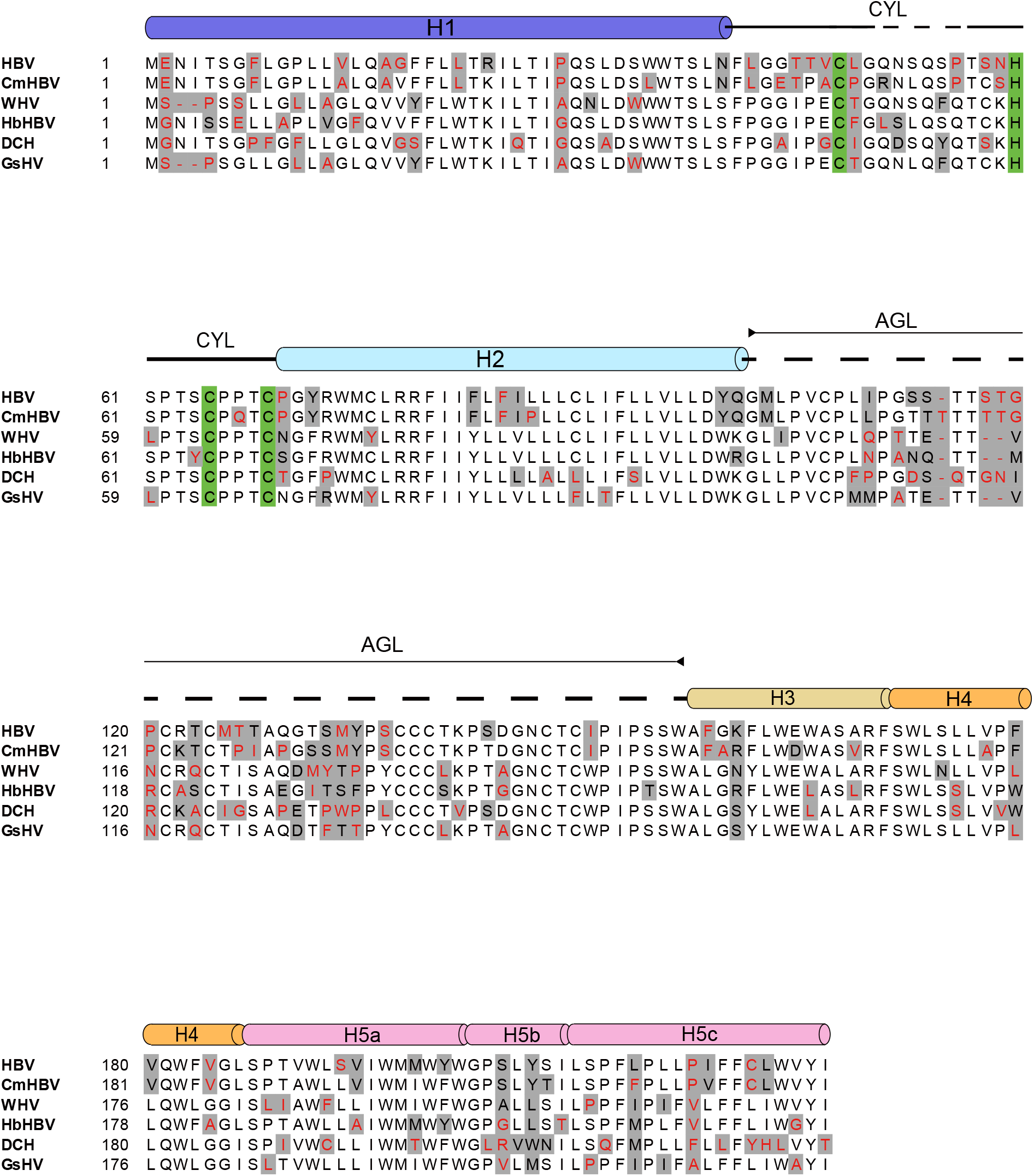
Sequence alignment of the surface antigens of several HBV-related viruses. Multiple sequence alignment (MSA) of S-HBsAg sequences of Hepatitis B virus (HBV, RefSeq: YP_009173871.1), Capuchin monkey hepatitis B virus (CmHBV, RefSeq: YP_009666526.1), Woodchuck hepatitis virus (WHV, RefSeq: NP_944491.1), Horseshoe bat hepatitis B virus (HbHBV, RefSeq: YP_009045996.1), Domestic cat hepadnavirus (DCH, RefSeq: YP_009553237.1) and Ground squirrel hepatitis virus (GsHV, RefSeq: NP_955537.1). The sequences are downloaded from NCBI. Helices are shown as cylinders. Residues which form the intracellular zinc finger motif on CYL are highlighted in green. Similar mutated residues were indicated in grey and the non-conserved residues were highlighted with red letters. Sequence alignment was performed with ClustalW and illustrated by BioEdit.

**Fig. S5.**
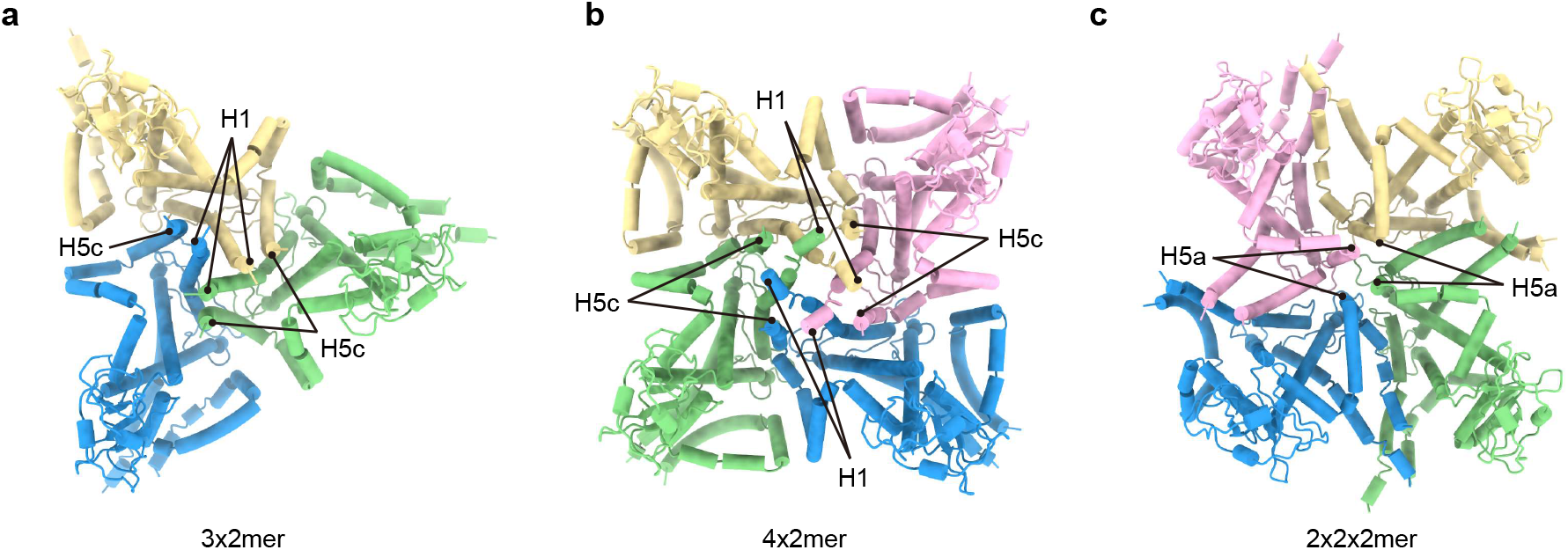
Assembly of HBsAg dimer in spherical SVP. **a-c**, Interactions between HBsAg dimer in the assembly of 3×2mer, 4×2mer, and 2×2×2mer respectively. Key helices involved in the assembly are indicated.

**Table S1.**
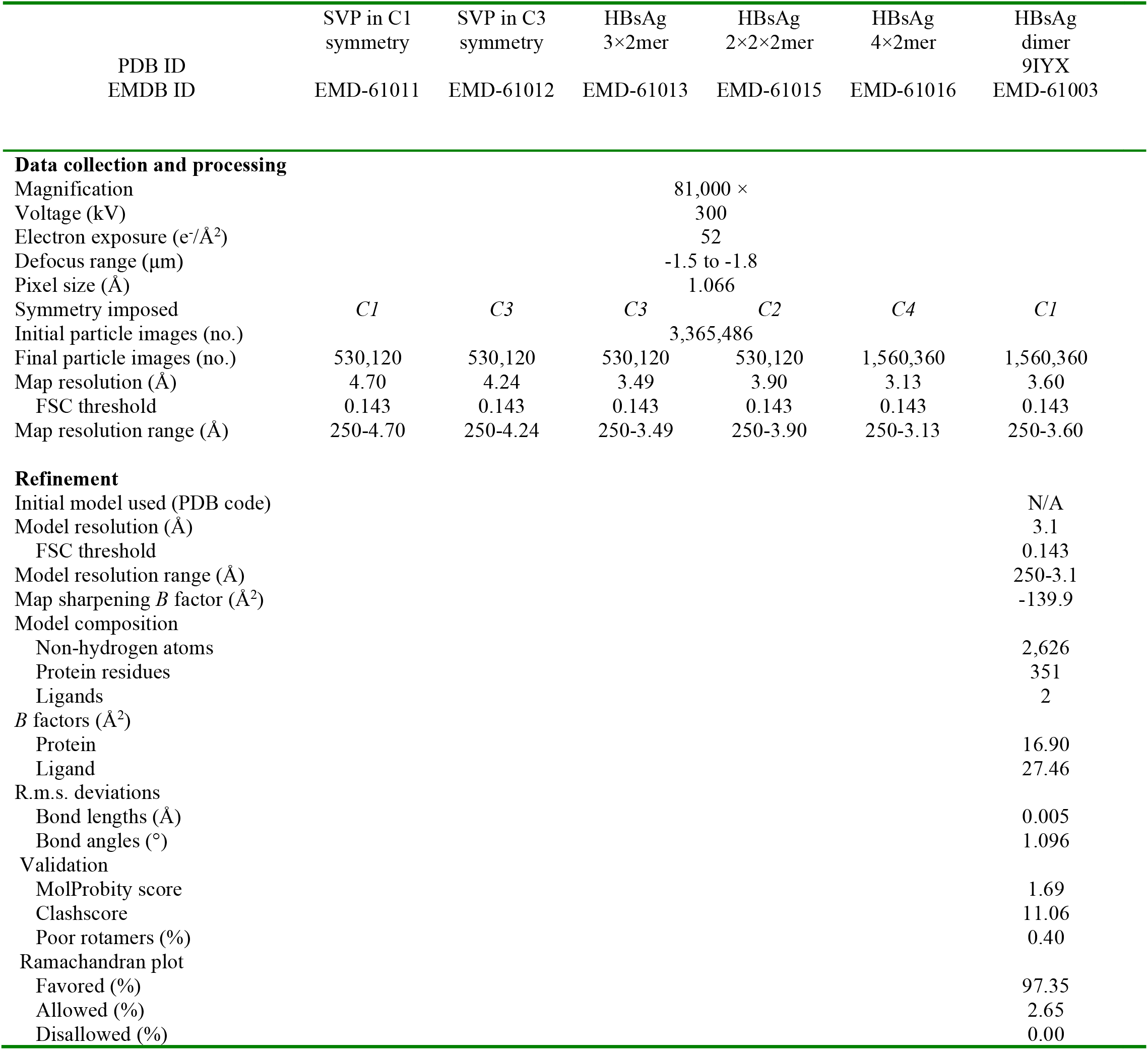
Cryo-EM data collection, refinement, and validation statistics. The parameters for the Cryo-EM data collection, processing, and validation of HBV SVP are listed in the table.

